# Largest Complete Mitochondrial Genome of a Gymnosperm, Sitka Spruce (*Picea sitchensis*), Indicates Complex Physical Structure

**DOI:** 10.1101/601104

**Authors:** Shaun D. Jackman, Lauren Coombe, René L. Warren, Heather Kirk, Eva Trinh, Tina McLeod, Stephen Pleasance, Pawan Pandoh, Yongjun Zhao, Robin J. Coope, Jean Bousquet, Joerg Bohlmann, Steven J. M. Jones, Inanc Birol

**Affiliations:** Genome Sciences Centre, BC Cancer, Vancouver, BC V5Z 4S6, Canada; Canada Research Chair in Forest Genomics, Institute for Sytems and Integrative Biology, Université Laval, Quebec, Quebec G1V 0A6, Canada; University of British Columbia, Michael Smith Laboratories, Vancouver, BC V6T 1Z4, Canada

## Abstract

Plant mitochondrial genomes vary widely in size. Although many plant mitochondrial genomes have been sequenced and assembled, the vast majority are of angiosperms, and few are of gymnosperms. Most plant mitochondrial genomes are smaller than a megabase, with a few notable exceptions. We have sequenced and assembled the 5.5 Mbp mitochondrial genome of Sitka spruce (*Picea sitchensis*), the largest complete mitochondrial genome of a gymnosperm. We sequenced the whole genome using Oxford Nanopore MinION, and then identified contigs of mitochondrial origin assembled from these long reads. The assembly graph shows a multipartite genome structure, composed of one smaller 168 kbp circular segment of DNA, and a larger 5.4 Mbp component with a branching structure. The assembly graph gives insight into a putative complex physical genome structure, and its branching points may represent active sites of recombination.

## Introduction

Plant mitochondrial genomes are amazingly diverse and complex (Mower et al. 2012). Land plant mitochondrial genomes range in size from 100 kilobases (kbp) for mosses such as *Mielichhoferia elongata* (Goruynov et al. 2018) to more than 11 megabases (Mbp) in the case of the flowering plant *Silene conica* (Sloan et al. 2012). Although their genome structure is often portrayed as a circle, the true physical structure of their genome appears to be a variety of circles, linear molecules, and complex branching structures (Backert et al. 1997; Backert & Börner 2000). While many species have a single master circle representation of their mitochondrial genome, others are composed of more than a hundred circular chromosomes (Sloan et al. 2012). The precise mechanism of how plant mitochondria replicate and maintain their DNA is not yet fully understood (Cupp & Nielsen 2014). It is hypothesized that recombination-dependent replication plays a role, giving a functional role to the repeat sequences often observed in mitochondria (Gualberto et al. 2014). This model does not fully explain how genomic copy number is regulated and maintained (Oldenburg & Bendich 2015), particularly in multipartite genomes (Vlcek et al. 2010). Although angiosperm mitochondrial genomes are well studied with numerous complete genomes available, few gymnosperm mitochondrial genomes are available: one from each of the cycads (Chaw et al. 2008), ginkgos, gnetales (Guo et al. 2016), and conifers (Jackman et al. 2015). Whereas other gymnosperm mitochondrial genomes are smaller than a megabase, conifer mitochondrial genomes can exceed five megabases (Jackman et al. 2015), larger than many bacteria. The origin and mechanism of this expansion are not known, but the trend correlates well with the very large nuclear genomes of conifers (De La Torre et al. 2014) and spruces in particular (Birol et al. 2013; Nystedt et al. 2013; Warren et al. 2015), compared to other plants. As plant mitochondrial genomes typically have fewer than 100 genes, what role this expanse of DNA serves, if any, remains mysterious.

Assembling plant mitochondrial genomes is difficult due to the presence of large (up to 30 kbp) perfect repeats, which may be involved in active recombination, and hypothesized recombination-dependent replication (Gualberto et al. 2014). A hybrid assembly of both long reads, which are able to span most repeats, and accurate short sequencing reads, which correct indel errors, is well suited to tackle these challenging genome features. When analyzing whole genome sequencing reads to reconstruct mitochondrial genomes, long reads provide additional confidence that the assembled sequences represent the true mitochondrial sequence in regions that share sequence homology between the mitochondrion, plastid, and nuclear genomes, due to the transfer of DNA between cellular compartments (Adams 2003; Smith 2011). Although hybrid assembly has been applied to assemble the plastid genome of *Eucalyptus pauciflora* (Wang et al. 2018), it has not yet been applied to the assembly of a plant mitochondrial genome.

Annotating plant mitochondrial genomes is also challenging, due to numerous features of plant mitochondria that are not typical of most organisms. For one, RNA editing of C-to-U is pervasive, and this process creates AUG start codons by editing ACG to AUG (Hiesel et al. 1989) or by editing GCG to GUG, an alternative start codon used by some plant mitochondrial genes (Sakamoto et al. 1997). RNA editing can also create stop codons in a similar fashion. Further complicating annotation using available bioinformatics pipelines, the typical GU-AG splice site expected by most splice-aware alignments tools is instead GNGCG-AY (Y denotes C or T) for group II introns (Lambowitz & Zimmerly 2010 and see results). Also, trans-spliced genes are common in mitochondrial genomes (Kamikawa et al. 2016), and no purpose-built software tool exists for identifying and annotating trans-spliced genes. To add further difficulty, trans-spliced exons may be as small as 22 bp, as is *nad5* exon 3 of gymnosperms (Guo et al. 2016) and other vascular plant mitochondria (Knoop et al. 1991). For these reasons, annotating a plant mitochondrial genome remains a laborious and manual task.

In this study, we report on the sequencing and assembly of the mitochondrial genome of Sitka spruce *(Picea sitchensis,* Pinaceae), a widely distributed conifer in the coastal regions of the Pacific Northwest. We show that this mitochondrial genome is one of the largest among plants and exhibits a multipartite genome structure.

## Methods

### Genome Sequencing and Assembly

Genomic DNA was extracted from young Sitka spruce (*Picea sitchensis* [Bong.] Carrière, genotype Q903) needles, as described in Coombe et al. (2016). We constructed 18 Oxford Nanopore 1D sequencing libraries, 16 by ligation of 1 to 7 μg of lightly needle-sheared genomic DNA and 2 by rapid transposition of 0.6 μg of unsheared genomic DNA, and sequenced these on 18 MinION R9.4 flow cells. This whole genome sequencing produced 98 Gbp in 9.6 million reads (SRA accession SRX5081713), yielding 5-fold depth of coverage of the roughly 20 Gbp nuclear genome, and 26-fold depth of coverage of the mitochondrial genome. Because separating putative mitochondrial reads by homology to known mitochondrial sequences could discard mitochondrial sequences that are unique to Sitka spruce, we chose to first assemble the whole genome reads and then compare contigs to known mitochondrial sequences. Assembling such a large number of Nanopore reads is not yet straight forward, and so we adopted an iterative approach to assembly. We first obtained a rough but computationally efficient assembly using Miniasm (Li 2016), after trimming adapter sequences with Porechop (Wick et al. 2017a). Miniasm produces an assembly whose sequencing error rate is comparable to that of the original reads, but no better. We polished this assembly using Racon (Vaser et al. 2017). We selected contigs with homology to the white spruce *(Picea glauca,* interior white spruce genotype PG29) mitochondrial genome (Jackman et al. 2015) using Bandage (Wick et al. 2015), retaining contigs with at least one 5 kbp alignment to the white spruce mitochondrion by BLASTN (Altschul et al. 1990). We note that this selection process would also discard any mitochondrial plasmids smaller than 5 kbp, if present.

We selected putative mitochondrial reads by aligning the Nanopore reads to this assembly using Minimap2 (Li 2018), and retained reads with an alignment score of 5000 or more. We assembled these reads using Unicycler (Wick et al. 2017b). This assembly yielded one circular contig and many linear contigs with no adjacent contigs, indicating that the assembly may not yet be complete, unless the genome were composed of linear chromosomes. We repeated the alignment of the Nanopore reads to the assembly, and again retained reads with an alignment score of 5000 or more. We assembled these reads using Flye (Kolmogorov et al. 2018), taking the output assembly graph `2-repeat/graph_final.gfa` that identifies repeats that are longer than the read length and determines their precise boundaries. This Flye assembly was polished using Racon. Contigs with homology to the white spruce mitochondrion were selected using Bandage, which uses BLASTN, requiring an alignment length of at least 5 kbp and percent identity at least 90. Contigs with unambiguous adjacent contigs were merged using the Bandage operation “Merge all possible nodes”.

In addition to a generally high sequencing error rate, Nanopore reads are particularly poor in accurately representing the length of homopolymer repeats. To compensate, we polished the assembly using one flow cell of Illumina HiSeq sequencing reads of the same DNA extraction, yielding 59-fold depth of coverage of the mitochondrial genome, to correct for sequencing and homopolymer length errors. We used Unicycler Polish to iteratively align the reads to the assembly using Bowtie2 (Langmead & Salzberg 2012), and correct the consensus sequence using Pilon (Walker et al. 2014). This iterative polishing process yielded no further corrections on the tenth round. Unicycler Polish applies Assembly Likelihood Estimate (ALE) (Clark et al. 2013) to each round to verify that the assembly of the final round of polishing resembles the reads the most. While annotating the genome, we found five indel errors in homopolymer runs that disrupted the reading frame of a gene. These five indel errors were corrected manually after inspecting the sequencing data.

### Annotation

We annotated coding genes and non-coding rRNA and tRNA genes using automated methods where possible, and performed manual inspection to refine these automated annotations. We used Prokka (Seemann 2014), which uses Prodigal (Hyatt et al. 2010) to identify single-exon coding genes and open reading frames (ORFs). We used MAKER (Holt & Yandell 2011), which uses BLASTP and Exonerate (Slater & Birney 2005) to identify cis-spliced coding genes. We used tRNAscan-SE (Lowe & Eddy 1997) and Aragorn (Laslett 2004) to identify tRNA. We used RNAmmer (Lagesen et al. 2007) and Barrnap (Seemann 2014) to identify rRNA. We used RNAweasel (Lang et al. 2007) and Infernal (Nawrocki et al. 2009) to identify group II introns. RFAM motif Domain-V (RM00007) represents domain V of group II introns, and RFAM family Intron_gpII (RF00029) represents both domains V and VI (Kalvari et al. 2017).

Following automated annotation, we reviewed coding genes for completeness, compared to their best BLASTP match, and corrected the annotation, most often for aspects that are particular to plant mitochondria. We manually corrected the annotation of genes to address start codons created by RNA editing of ACG to the start codon AUG, and editing of GCG to the alternative start codon GUG (see results for details). Three genes display atypical start codons: *rpl16* uses a GUG start codon (Sakamoto et al. 1997); *rps19* uses a GUG start codon created by RNA editing GCG, seen also in *Pinus strobus* AJP33554.1; *matR* appears to use an unusual GGG start codon, seen also in *Cycas taitun-gensis* YP_001661429.1 (Chaw et al. 2008) and *Pinus strobus* AJP33535.1. The gene *sdh4* was missed by automatic annotation, as its coding sequence was found to overlap with *cox3* by 73 bp on the same strand.

We reviewed splice sites, and adjusted their position to agree with the expected splicing motifs of group II introns when possible, ensuring not to introduce insertions or deletions into the peptide sequence compared to homologous proteins. We confirmed the presence of domain V of the group II intron upstream of the 3’ splice site, identified by RNAweasel or Infernal. We manually annotated trans-spliced introns by comparing alignments of homologous proteins to the genome. We determined the 5’ and 3’ splice sites similarly to cis-spliced introns, looking for expected group II splicing motifs, and domain V upstream of the 3’ splice site. When Infernal did find a match to RFAM Intron_gpII (RF00029), it frequently identified the precise 3’ splice site, in agreement with protein sequence homology.

The scripts to assemble and annotate the Sitka spruce mitochondrial genome are available online at https://github.com/sjackman/psitchensismt.

## Results and Discussion

### Complete Genome Assembly

The complete mitochondrial genome of Sitka spruce is 5.52 Mbp assembled in 13 segments, and has a GC content of 44.7%. The genome assembly is composed of two components: a 168 kbp circular segment, and a larger 5.36 Mbp component composed of 12 segments, visualized by Bandage (Wick et al. 2015) in Fig. 1. The two smallest segments (27 kbp and 24 kbp, labeled 12 and 13 respectively) exhibit an estimated copy number of two based on their depth of sequencing coverage, and all other segments have similar depth of coverage, assumed to represent single copy. The single-copy segments range in size from 84 kbp to 1.65 Mbp. No sequence variation is evident in these repeats. An absence of variation in the repeat implies that they may be involved in active recombination (Maréchal & Brisson 2010). Though 10% of reads are larger than 24 kbp, no reads fully span these repeats.

**Figure 1:**
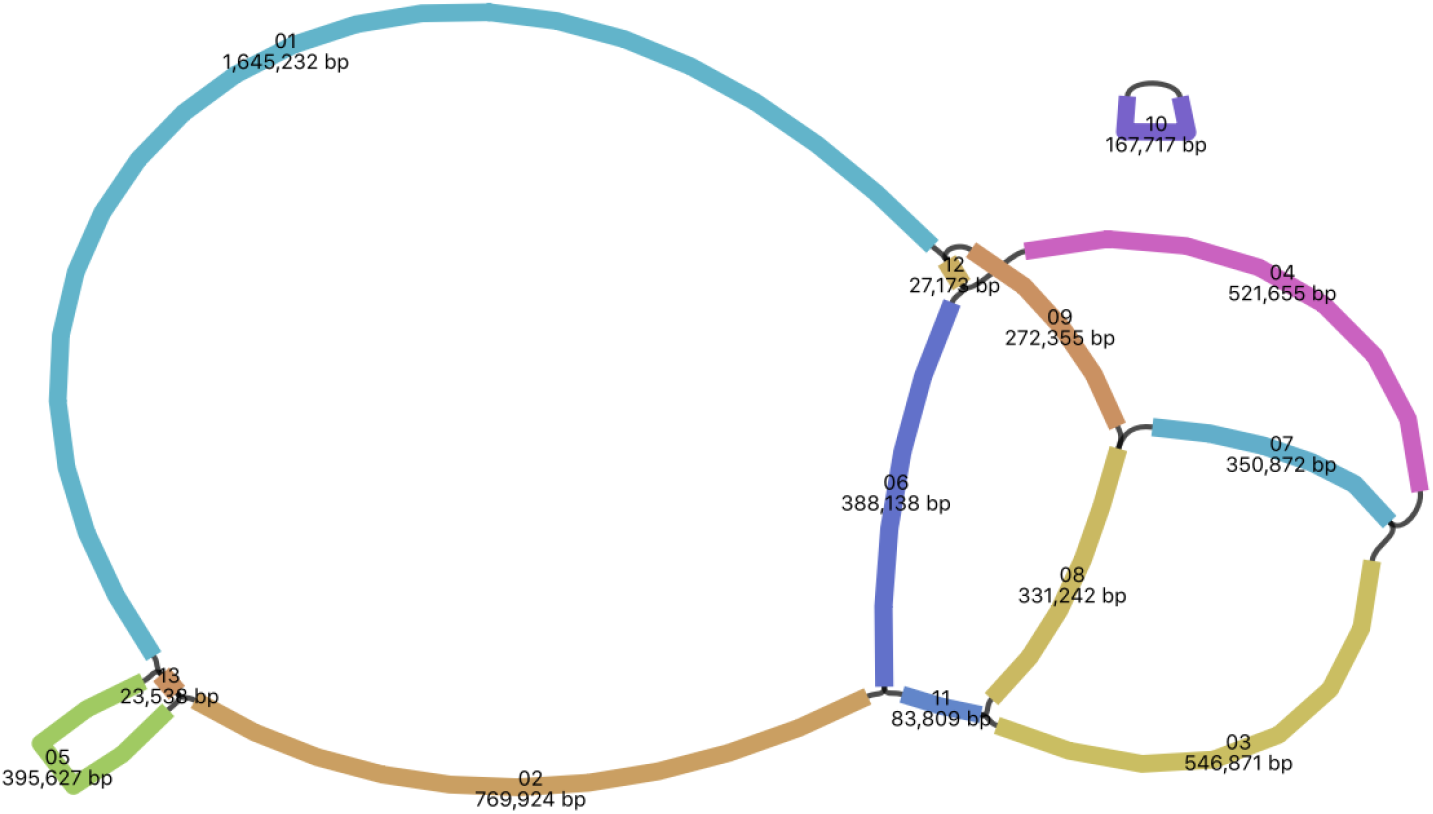
The assembly graph of the mitochondrial genome of Sitka spruce. Each segment is labeled with its size and named 01 through 13 by rank of size.

The draft mitochondrial genome assembly of white spruce (Jackman et al. 2015) was assembled from paired-end and mate-pair Illumina sequencing. The draft assembly is composed of 117 contigs larger than 2 kbp arranged in 36 scaffolds with a contig N50 of 102 kbp and scaffold N50 of 369 kbp. It provided little information as to the structure of the mitochondrial genome. The complete mitochondrial genome assembly of Sitka spruce assembled from Oxford Nanopore sequencing is composed of 13 contigs larger than 20 kbp with a N50 of 547 kbp. The assembly graph (Fig. 1) reveals a multipartite genome structure. The use of long reads were critical in achieving this contiguity and completeness.

The complete genome is composed of 1.7% (93 kbp) of genes with known function, 28.0% (1,545 kbp) of 6,806 ORFs (each of at least 90 bp), 3.7% (205 kbp) of repeats, and 66.6% unclassified sequences. Of the ORFs, 1,039 are at least 300 bp (100 amino acids) in size and compose 7.2% (400 kbp) of the genome. Aligning the ORFs with BLASTP, 63 ORFs (17 ORFs of at least 300 bp) have a significant (E < 0.001) hit to the nr database. Plastid-derived sequences composes 0.25% (14 kbp) of the genome spread across 24 segments.

The nuclear repeats LTR/Gypsy compose 51% of the repeat sequence, LTR/Copia compose 7%, simple repeat sequences compose 34%, low complexity compose 3%, and 5% are other repeat sequences. The 36-bp Bpu repeat sequence is present in roughly 500 copies in *Cycas taitungensis* and roughly 100 copies in *Ginkgo biloba* (Guo et al. 2016). We find only a single full-length copy with four mismatches in Sitka spruce, similar to *Welwitschia mirabilis.*

### Genes

The mitochondrial genome of Sitka spruce has 41 distinct protein coding genes with known function, 3 distinct rRNA genes (Table 1), and 27 distinct tRNA genes representing 18 distinct anticodons (Table 2). The relative order and orientation of these genes are shown in Fig. 2. The 41 known protein coding genes found in the gymnosperm mitochondria *Cycas taitungensis* (Chaw et al. 2008) and *Ginkgo biloba* (Guo et al. 2016) are also found in Sitka spruce. The 29 introns, 16 cis-spliced and 13 trans-spliced, are found in 10 protein coding genes, two pseudogenes, and one plastid-derived tRNA (Table 3).

**Figure 2:**
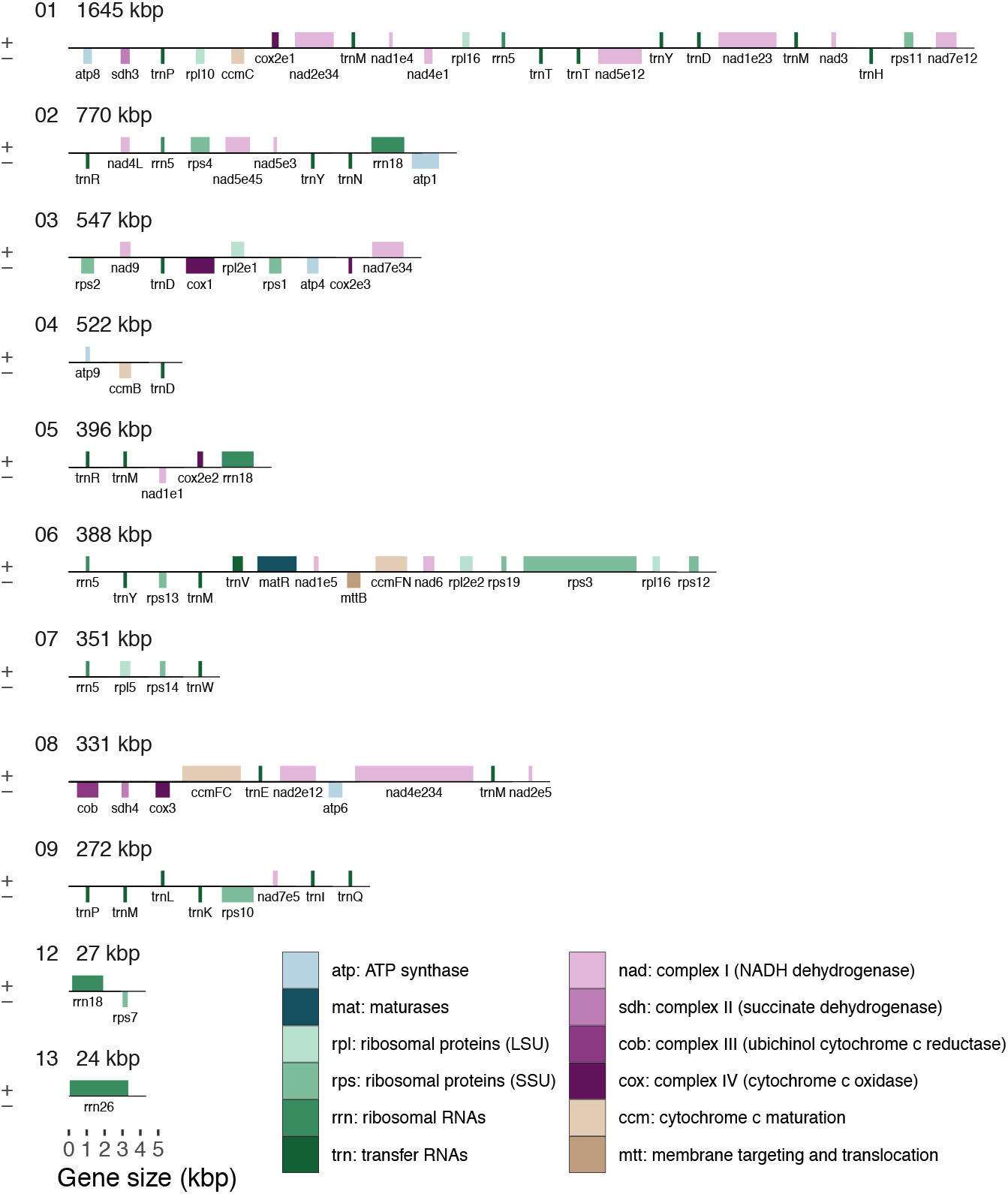
The order, orientation, and size of the genes of Sitka spruce. Each box is proportional to the size of the gene including introns, except that genes smaller than 200 nucleotides are shown as 200 nucleotides. Intergenic regions are not to scale.

**Table 1:** rRNA gene content of four gymnosperms. This table is adapted from Table S1 of Guo et al. (2016) with the addition of Sitka spruce. *One copy is present on a repeat segment with an estimated copy number of two.

| Gene | Cycas | Ginkgo | Sitka | Welwitschia |
| --- | --- | --- | --- | --- |
| *rrn5* | 1 | 1 | 4 | 1 |
| *rrn18* | 1 | 1 | 3* | 1 |
| *rrn26* | 1 | 1 | 1* | 1 |
| **Total rRNA** | **3** | **3** | **8** | **3** |

**Table 2:** tRNA content of four gymnosperms. Sitka spruce has 27 tRNA genes, one of which is cis-spliced, with 18 distinct anticodons, coding for 15 distinct amino acids. This table is adapted from Table S1 of Guo et al. (2016) with the addition of Sitka spruce. (i) Contains a cis-spliced group II intron. *Anticodon is inferred to be edited (Weber et al. 1990).

| Gene | Cycas | Ginkgo | Sitka | Welwitschia |
| --- | --- | --- | --- | --- |
| trnC-GCA | 1 | 1 | - | - |
| trnD-GUC | 1 | 1 | 3 | 1 |
| trnE-UUC | 1 | 1 | 1 | 1 |
| trnF-GAA | 1 | 2 | - | - |
| trnG-GCC | 1 | - | - | - |
| trnG-UCC | - | 1* | - | - |
| trnH-GUG | 1 | 1 | 1 | - |
| trnI-CAU | 1* | 1* | 1* | 1* |
| trnK-UUU | 1 | 1* | 1 | - |
| trnL-CAA | - | - | 1 | - |
| trnL-UAA | 1* | 2 | - | - |
| trnL-UAG | 1 | 1 | - | - |
| trnM-CAU | 6 | 2 | 6 | 1 |
| trnN-GUU | 1 | - | 1 | - |
| trnP-AGG | 1 | 1 | 1 | - |
| trnP-UGG | 1 | 1 | 1 | - |
| trnQ-UUG | 1* | 1 | 1 | 1 |
| trnR-ACG | - | - | - | 1 |
| trnR-CCG | - | - | 1 | - |
| trnR-GCG | - | - | 1 | - |
| trnR-UCU | 1* | 1 | - | - |
| trnS-GCU | 1 | 1 | - | - |
| trnS-GGA | 1 | - | - | - |
| trnS-UGA | 1 | 1 | - | - |
| trnT-AGU | - | - | 1 | - |
| trnT-UGU | - | - | 1 | - |
| trnV-UAC (i) | 1 | - | 1 | - |
| trnW-CCA | 1 | 2 | 1 | 1 |
| trnY-AUA | - | - | 1 | - |
| trnY-GUA | 1 | 1 | 2 | 1 |
| **Total tRNA** | **27** | **23** | **27** | **8** |

**Table 3:**
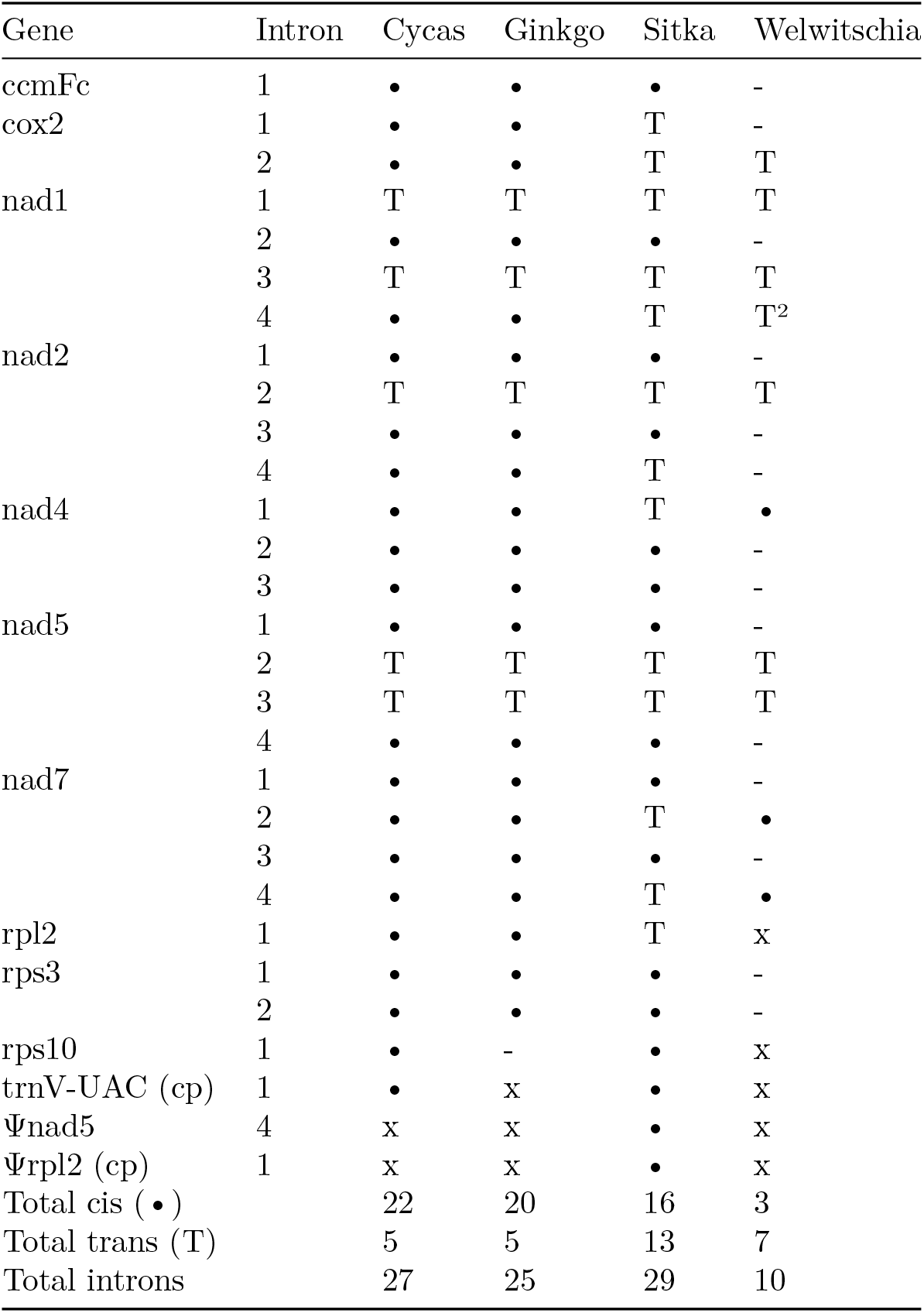
Intron content of four gymnosperms. Sitka spruce has 29 introns, 16 cis-spliced (•) and 13 trans-spliced (T), in ten protein coding genes, two pseudogenes (Ψ), and one tRNA. “T^2^” indicates a tripartite (double trans-spliced) intron. “-” indicates intron absence. “x” indicates gene absence. “cp” indicates plastid-derived. This table is adapted from Guo et al. (2016) with the addition of Sitka spruce.

| Gene | Intron | Cycas | Ginkgo | Sitka | Welwitschia |
| --- | --- | --- | --- | --- | --- |
| ccmFc | 1 | • | • | • | - |
| cox2 | 1 | • | • | T | - |
|  | 2 | • | • | T | T |
| nad1 | 1 | T | T | T | T |
|  | 2 | • | • | • | - |
|  | 3 | T | T | T | T |
|  | 4 | • | • | T | T<sup>2</sup> |
| nad2 | 1 | • | • | • | - |
|  | 2 | T | T | T | T |
|  | 3 | • | • | • | - |
|  | 4 | • | • | T | - |
| nad4 | 1 | • | • | T | • |
|  | 2 | • | • | • | - |
|  | 3 | • | • | • | - |
| nad5 | 1 | • | • | • | - |
|  | 2 | T | T | T | T |
|  | 3 | T | T | T | T |
|  | 4 | • | • | • | - |
| nad7 | 1 | • | • | • | - |
|  | 2 | • | • | T | • |
|  | 3 | • | • | • | - |
|  | 4 | • | • | T | • |
| rpl2 | 1 | • | • | T | x |
| rps3 | 1 | • | • | • | - |
|  | 2 | • | • | • | - |
| rps10 | 1 | • | - | • | x |
| trnV-UAC (cp) | 1 | • | x | • | x |
| Ψnad5 | 4 | x | x | • | x |
| Ψrpl2 (cp) | 1 | x | x | • | x |
| Total cis (•) |  | 22 | 20 | 16 | 3 |
| Total trans (T) |  | 5 | 5 | 13 | 7 |
| Total introns |  | 27 | 25 | 29 | 10 |

Four ORFs (937, 507, 371, and 221 amino acids) contain an organellar DNA polymerase type B (DNA_pol_B_2) Pfam family (El-Gebali et al. 2018). The largest one also contains a segment homologous to the structural maintenance of chromosome (SMC_N) Pfam family, which includes the *RecF* and *RecN* proteins involved in DNA recombination. We hypothesize that this ORF may be involved in recombination-dependent replication of mitochondrial DNA (Gualberto et al. 2014). These ORFs have homology to DNA polymerase genes found in the mitochondria of *Picea glauca* and the angiosperms *Cocos nucifera, Daucus carota, Helianthus annuus,* and *Silene vulgaris.* Two ORFs (781 and 560 amino acids) contain an RNA polymerase (RNA_pol) domain. These ORFs have homology to DNA-dependent RNA polymerase genes found in the mitochondria of *Picea glauca* and the angiosperms *Beta vulgaris, Cocos nucifera, Daucus carota,* and *Phoenix dactylifera.* The two largest genes of the Sitka spruce mitochondrial genome are these putative DNA and RNA polymerase genes.

The *matR* gene and three additional ORFs (476, 197, and 163 amino acids) contain a reverse transcriptase, or RNA-dependent DNA polymerase, (RT_like) NCBI conserved protein domain with similarity to the group II intron reverse transcriptase/maturase (group_II_RT_mat) NCBI conserved protein domain (Marchler-Bauer et al. 2016). We hypothesize that these ORFs may be additional maturases involved in splicing (Matsuura 2001). These ORFs have homology to mitochondrial genes of *Picea glauca* and the angiosperm *Utricularia reniformis.*

The full complement of rRNA genes are present in Sitka spruce, shown in Table 1. Unlike rRNA genes of other gymnosperms, the Sitka spruce rRNA genes are present in multiple copies. The 5S rRNA gene *rrn5* is present in four copies. The small subunit rRNA gene *rrn18* is present in three copies, though one copy is found on the 27 kbp repeat segment with an estimated copy number of two. One copy of the large subunit rRNA gene *rrn26* is present, though it is found on the 24 kbp repeat segment, which also has an estimated copy number of two.

Sitka spruce has 27 tRNA genes, representing 18 distinct anticodons, coding for 15 distinct amino acids, DEHIKLMNPQRTVWY (Table 2). tRNA genes coding for the amino acids ACFGS are absent in Sitka spruce, and also absent in *Welwitschia. trnM-CAU* exhibits six copies, *trnD-GUC* three copies, and *trnY-GUA* two copies. All other tRNA genes are single copy. *trnN-GUU, trnV-UAC,* and one copy of *trnfM-CAU* are derived from plastid origins. One cis-spliced intron is observed in the plastid-derived *trnV-UAC* gene, also seen in *Cycas taitungensis.* Six tRNA genes (*trnL-CAA, trnR-CCG, trnR-GCG, trnT-AGU, trnT-UGU,* and *trnY-AUA)* found in Sitka spruce are absent in *Cycas, Ginkgo,* and *Welwitschia*.

In addition to three plastid-derived tRNA genes, ten partial plastid genes are found in the 14 kbp of plastid-derived sequence: *atpB, atpE, atpF, chlN, petA, psaA, rps3, rrn18,* a partial copy of *rpl2,* and a partial *trnS-GGA* gene with homology to *Cycas taitungensis.* The *rpl2* partial gene is more similar to eudicot plastids (77% identical to *Helwingia himalaica, Robinia pseudoacacia,* and many other eudicots) than it is to the Sitka spruce plastid (66% identical).

### Introns

Although the same 27 introns are found in the same 11 genes as *Cycas taitungensis* (Chaw et al. 2008; Guo et al. 2016), eight introns that are cis-spliced in *Cycas* are trans-spliced in Sitka spruce, more than doubling the number of trans-spliced introns found in *Cycas.* Nearly half of the introns in Sitka spruce are trans-spliced. All introns are group II introns, whose domain V was identified by both RNAweasel and Infernal, with one exception.

The first intron of *nad1* is trans-spliced in Sitka spruce and other gymnosperms (Guo et al. 2016). No domain V is detectable by Infernal either downstream of exon 1 or upstream of exon 2 in Sitka spruce nor any of *Cycas, Ginkgo,* and *Welwitschia*. The genomic disruption of this intron may occur in domain V itself, as is seen in *cox2* of *Diphylleia rotans* (Kamikawa et al. 2016).

The fourth intron of *nad1* is cis-spliced, and it contains *matR* in *Cycas,* trans-spliced with a single disruption in Sitka spruce, and trans-spliced with two distinct genomic disruptions in *Welwitschia mirabilis* (Figure S2 of Guo et al. 2016). Whereas *matR* is found in a cis-spliced intron in *Cycas* and free-standing in *Welwitschia*, it is found upstream of *nad1* exon 5 in Sitka spruce. In this regard, Sitka spruce appears to be an evolutionary midpoint between *Cycas* and *Welwitschia*. Sitka spruce has not however experienced the extensive gene loss observed in *Welwitschia*.

A second partial copy of *nad5* is found in Sitka spruce with one cis-spliced group II intron, representing exons 4 and 5. The translated protein sequence of this partial gene is more similar however to eudicot mitochondria (99% identical to both *Chrysobalanus icaco* and *Hirtella racemosa,* >95% identical to many other eudicots, and 94% identical to one monocot *Triantha glutinosa)* than to the complete *nad5* of Sitka spruce (76% identical). It may have been acquired by horizontal gene transfer, as is frequently reported in plant mitochondria (Richardson & Palmer 2006) of both gymnosperms (Won & Renner 2003) and angiosperms (Bergthorsson et al. 2003). This interpretation of horizontal gene transfer in plant mitochondria is not universally accepted (Goremykin et al. 2008). This partial copy of *nad5* is also found in white spruce (Jackman et al. 2015) with 100% nucleotide identity. This level of conservation between Sitka spruce, white spruce, and angiosperms suggests that this partial gene may be functional. We find no upstream domain V, whose presence would indicate that it may be part of a larger trans-spliced gene. A putatitve alternative GUG start codon created by RNA editing of GCG could initiate translation of this partial gene.

RNAweasel identifies 34 group II domain V regions in Sitka spruce, 26 of which are associated with the intron of a gene. Two domain V regions are found in the cis-spliced introns of the pseudogenes *Ψnad5* and plastid-derived *Ψrpl2.* The remaining six domain V regions are not associated with a gene, and further investigation would be needed to determine whether they may also be partial fragments of pseudogene introns.

The splice-site motifs of the 14 cis-spliced genes of the Sitka spruce mitochondrial genome are shown in Fig. 3, visualized by WebLogo (Crooks 2004). Because its position is variable, the bulged adenosine of the 3’ splice site, typically found at position −7 or −8, is not readily apparent.

**Figure 3:**
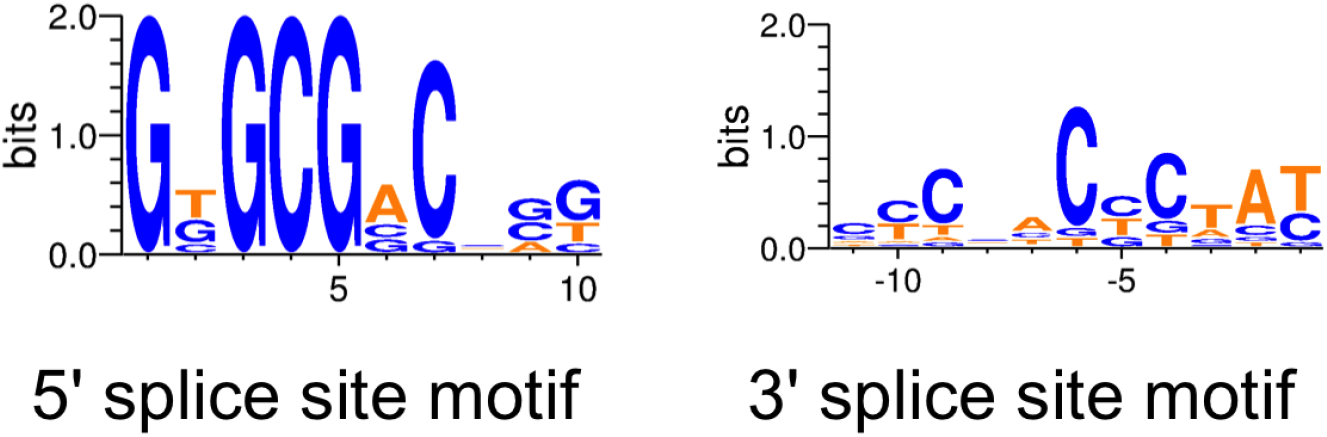
The 5’ and 3’ splice-site motifs of the 14 cis-spliced genes of Sitka spruce.

## Conclusion

The 5.5 Mbp mitochondrial genome of Sitka spruce is among the largest ones in plants, and is the largest complete mitochondrial genome reported for a gymnosperm. It follows the trend seen for spruce and conifer nuclear genomes, which are also among the largest in plants (De La Torre et al. 2014). The physical structure of the Sitka spruce mitochondrial genome is not the typical circularly-mapping single chromosome, but multipartite. The larger component of the assembly graph exhibits a rosette-like structure, mirroring the rosette-like structures observed in electron micrographs of mitochondrial DNA (Backert & Börner 2000). The intricate structure and the conservation of large repeat elements suggest the presence of active sites for hypothesized recombination-dependent replication of the mitochondrial genome (Gualberto et al. 2014). Considering heteroplasmy resulting from naturally hybridizing species of spruce and paternal leakage of mitochondria, intermolecular recombination would result in interspecific hybridization of the mitochondrial genome, which has been reported to occur in natural spruce populations (Jaramillo-Correa & Bousquet 2005). Although sequence identity of the set of common mitochondrial proteins is well conserved, their splicing structure is not. Trans-splicing is a frequently-employed mechanism of plant mitochondria to compensate for genomic structural instability, and Sitka spruce has a record number of trans-spliced introns. As the first long read assembly of a complete plant mitochondrial genome, also exhibiting a multipartite genome structure, this resource should prove invaluable to future investigations into the genome structure and mechanism of replication of conifer mitochondrial genomes.

## Data Availability

The Oxford Nanopore MinION sequencing data has SRA accessions SRX5081713, and SRR8264752 through SRR8264769. The Illumina HiSeq sequencing data has SRA accession SRR5028199. The Sitka spruce *(Picea sitchensis*) genotype Q903 complete mitochondrial genome has NCBI GenBank accessions MK697696 through MK697708.

## Acknowledgements

Funding was provided by Genome Canada, Genome Quebec, Genome British Columbia, and Genome Alberta for the Spruce-Up (243FOR) Project (http://www.spruce-up.ca).

## Author Contributions

SDJ drafted the manuscript. SDJ, LC, RLW, SP, JoB, and IB revised the manuscript. HK, ET, and TM constructed the libraries and sequenced the DNA. SP, PP, YZ, and RC supervised the sequencing. SDJ assembled and annotated the mitochondrial genome. SDJ designed and executed the data analysis. JeB, JoB, SJMJ, and IB supervised the project.

## Competing Interests

The authors declare no competing interests.

## References

Adams K. 2003. Evolution of mitochondrial gene content: gene loss and transfer to the nucleus. Molecular Phylogenetics and Evolution. 29:380–395. doi: 10.1016/s1055-7903(03)00194-5.

Altschul SF, Gish W, Miller W, Myers EW, Lipman DJ. 1990. Basic local alignment search tool. Journal of Molecular Biology. 215:403–410. doi: 10.1016/s0022-2836(05)80360-2.

Backert S, Börner T. 2000. Phage T4-like intermediates of DNA replication and recombination in the mitochondria of the higher plant Chenopodium album (L.). Current Genetics. 37:304–314. doi: 10.1007/s002940050532.

Backert S, Lynn Nielsen B, Börner T. 1997. The mystery of the rings: structure and replication of mitochondrial genomes from higher plants. Trends in Plant Science. 2:477–483. doi: 10.1016/s1360-1385(97)01148-5.

Bergthorsson U, Adams KL, Thomason B, Palmer JD. 2003. Widespread horizontal transfer of mitochondrial genes in flowering plants. Nature. 424:197–201. doi: 10.1038/nature01743.

Birol I et al. 2013. Assembling the 20 Gb white spruce (Picea glauca) genome from whole-genome shotgun sequencing data. Bioinformatics. 29:1492–1497. doi: 10.1093/bioinformatics/btt178.

Chaw S-M et al. 2008. The Mitochondrial Genome of the Gymnosperm Cycas taitungensis Contains a Novel Family of Short Interspersed Elements, Bpu Sequences, and Abundant RNA Editing Sites. Molecular Biology and Evolution. 25:603–615. doi: 10.1093/molbev/msn009.

Clark SC, Egan R, Frazier PI, Wang Z. 2013. ALE: a generic assembly likelihood evaluation framework for assessing the accuracy of genome and metagenome assemblies. Bioinformatics. 29:435–443. doi: 10.1093/bioinformatics/bts723.

Coombe L et al. 2016. Assembly of the Complete Sitka Spruce Chloroplast Genome Using 10X Genomics’ GemCode Sequencing Data Budak, H, editor. PLOS ONE. 11:e0163059. doi: 10.1371/journal.pone.0163059.

Crooks GE. 2004. WebLogo: A Sequence Logo Generator. Genome Research. 14:1188–1190. doi: 10.1101/gr.849004.

Cupp JD, Nielsen BL. 2014. Minireview: DNA replication in plant mitochondria. Mitochondrion. 19:231–237. doi: 10.1016/j.mito.2014.03.008.

De La Torre AR et al. 2014. Insights into Conifer Giga-Genomes. PLANT PHYSIOLOGY. 166:1724–1732. doi: 10.1104/pp.114.248708.

El-Gebali S et al. 2018. The Pfam protein families database in 2019. Nucleic Acids Research. 47:D427–D432. doi: 10.1093/nar/gky995.

Goremykin VV, Salamini F, Velasco R, Viola R. 2008. Mitochondrial DNA of Vitis vinifera and the Issue of Rampant Horizontal Gene Transfer. Molecular Biology and Evolution. 26:99–110. doi: 10.1093/molbev/msn226.

Goruynov DV et al. 2018. Complete mitochondrial genome sequence of the ‘copper moss’ Mielichhoferia elongata reveals independent nad7 gene functionality loss. PeerJ. 6:e4350. doi: 10.7717/peerj.4350.

Gualberto JM et al. 2014. The plant mitochondrial genome: Dynamics and maintenance. Biochimie. 100:107–120. doi: 10.1016/j.biochi.2013.09.016.

Guo W et al. 2016. GinkgoandWelwitschiaMitogenomes Reveal Extreme Contrasts in Gymnosperm Mitochondrial Evolution. Molecular Biology and Evolution. 33:1448–1460. doi: 10.1093/molbev/msw024.

Hiesel R, Wissinger B, Schuster W, Brennicke A. 1989. RNA editing in plant mitochondria. Science. 246:1632–1634. doi: 10.1126/science.2480644.

Holt C, Yandell M. 2011. MAKER2: an annotation pipeline and genome-database management tool for second-generation genome projects. BMC Bioinformatics. 12. doi: 10.1186/1471-2105-12-491.

Hyatt D et al. 2010. Prodigal: prokaryotic gene recognition and translation initiation site identification. BMC Bioinformatics. 11. doi: 10.1186/1471-2105-11-119.

Jackman SD et al. 2015. Organellar Genomes of White Spruce (Picea glauca): Assembly and Annotation. Genome Biology and Evolution. 8:29–41. doi: 10.1093/gbe/evv244.

Jaramillo-Correa JP, Bousquet J. 2005. Mitochondrial Genome Recombination in the Zone of Contact Between Two Hybridizing Conifers. Genetics. 171:1951–1962. doi: 10.1534/genetics.105.042770.

Kalvari I et al. 2017. Rfam 13.0: shifting to a genome-centric resource for non-coding RNA families. Nucleic Acids Research. 46:D335–D342. doi: 10.1093/nar/gkx1038.

Kamikawa R, Shiratori T, Ishida K-I, Miyashita H, Roger AJ. 2016. Group II Intron-MediatedTrans-Splicing in the Gene-Rich Mitochondrial Genome of an Enigmatic Eukaryote, Diphylleia rotans. Genome Biology and Evolution. 8:458–466. doi: 10.1093/gbe/evw011.

Knoop V, Schuster W, Wissinger B, Brennicke A. 1991. Trans splicing integrates an exon of 22 nucleotides into the nad5 mRNA in higher plant mitochondria. The EMBO Journal. 10:3483–3493. doi: 10.1002/j.1460-2075.1991.tb04912.x.

Kolmogorov M, Yuan J, Lin Y, Pevzner P. 2018. Assembly of Long Error-Prone Reads Using Repeat Graphs. doi: 10.1101/247148.

Lagesen K et al. 2007. RNAmmer: consistent and rapid annotation of ribosomal RNA genes. Nucleic Acids Research. 35:3100–3108. doi: 10.1093/nar/gkm160.

Lambowitz AM, Zimmerly S. 2010. Group II Introns: Mobile Ribozymes that Invade DNA. Cold Spring Harbor Perspectives in Biology. 3:a003616–a003616. doi: 10.1101/cshperspect.a003616.

Lang BF, Laforest M-J, Burger G. 2007. Mitochondrial introns: a critical view. Trends in Genetics. 23:119–125. doi: 10.1016/j.tig.2007.01.006.

Langmead B, Salzberg SL. 2012. Fast gapped-read alignment with Bowtie 2. Nature Methods. 9:357–359. doi: 10.1038/nmeth.1923.

Laslett D. 2004. ARAGORN, a program to detect tRNA genes and tmRNA genes in nucleotide sequences. Nucleic Acids Research. 32:11–16. doi: 10.1093/nar/gkh152.

Li H. 2018. Minimap2: pairwise alignment for nucleotide sequences Birol, I, editor. Bioinformatics. 34:3094–3100. doi: 10.1093/bioinformatics/bty191.

Li H. 2016. Minimap and miniasm: fast mapping and de novo assembly for noisy long sequences. Bioinformatics. 32:2103–2110. doi: 10.1093/bioinformatics/btw152.

Lowe TM, Eddy SR. 1997. tRNAscan-SE: A Program for Improved Detection of Transfer RNA Genes in Genomic Sequence. Nucleic Acids Research. 25:0955–964. doi: 10.1093/nar/25.5.0955.

Marchler-Bauer A et al. 2016. CDD/SPARCLE: functional classification of proteins via subfamily domain architectures. Nucleic Acids Research. 45:D200–D203. doi: 10.1093/nar/gkw1129.

Maréchal A, Brisson N. 2010. Recombination and the maintenance of plant organelle genome stability. New Phytologist. 186:299–317. doi: 10.1111/j.1469-8137.2010.03195.x.

Matsuura M. 2001. Mechanism of maturase-promoted group II intron splicing. The EMBO Journal. 20:7259–7270. doi: 10.1093/emboj/20.24.7259.

Mower JP, Sloan DB, Alverson AJ. 2012. Plant Mitochondrial Genome Diversity: The Genomics Revolution. Plant Genome Diversity Volume 1. 123–144. doi: 10.1007/978-3-7091-1130-7_9.

Nawrocki EP, Kolbe DL, Eddy SR. 2009. Infernal 1.0: inference of RNA alignments. Bioinformatics. 25:1335–1337. doi: 10.1093/bioinformatics/btp157.

Nystedt B et al. 2013. The Norway spruce genome sequence and conifer genome evolution. Nature. 497:579–584. doi: 10.1038/nature12211.

Oldenburg DJ, Bendich AJ. 2015. DNA maintenance in plastids and mitochondria of plants. Frontiers in Plant Science. 6. doi: 10.3389/fpls.2015.00883.

Richardson AO, Palmer JD. 2006. Horizontal gene transfer in plants. Journal of Experimental Botany. 58:1–9. doi: 10.1093/jxb/erl148.

Sakamoto W, Tan S-H, Murata M, Motoyoshi F. 1997. An Unusual Mitochondrial atp9-rpl16 Cotranscript Found in the Maternal Distorted Leaf Mutant of Arabidopsis thaliana: Implication of GUG as an Initiation Codon in Plant Mitochondria. Plant and Cell Physiology. 38:975–979. doi: 10.1093/oxfordjournals.pcp.a029261.

Seemann T. 2014. Prokka: rapid prokaryotic genome annotation. Bioinformatics. 30:2068–2069. doi: 10.1093/bioinformatics/btu153.

Slater G, Birney E. 2005. BMC Bioinformatics. 6:31. doi: 10.1186/1471-2105-6-31.

Sloan DB et al. 2012. Rapid Evolution of Enormous, Multichromosomal Genomes in Flowering Plant Mitochondria with Exceptionally High Mutation Rates Gray, MW, editor. PLoS Biology. 10:e1001241. doi: 10.1371/journal.pbio.1001241.

Smith DR. 2011. Extending the Limited Transfer Window Hypothesis to Inter-organelle DNA Migration. Genome Biology and Evolution. 3:743–748. doi: 10.1093/gbe/evr068.

Vaser R, Sović I, Nagarajan N, Šikić M. 2017. Fast and accurate de novo genome assembly from long uncorrected reads. Genome Research. 27:737–746. doi: 10.1101/gr.214270.116.

Vlcek C, Marande W, Teijeiro S, Lukeš J, Burger G. 2010. Systematically fragmented genes in a multipartite mitochondrial genome. Nucleic Acids Research. 39:979–988. doi: 10.1093/nar/gkq883.

Walker BJ et al. 2014. Pilon: An Integrated Tool for Comprehensive Microbial Variant Detection and Genome Assembly Improvement Wang, J, editor. PLoS ONE. 9:e112963. doi: 10.1371/journal.pone.0112963.

Wang W et al. 2018. Assembly of chloroplast genomes with long- and short-read data: a comparison of approaches using Eucalyptus pauciflora as a test case. BMC Genomics. 19. doi: 10.1186/s12864-018-5348-8.

Warren RL et al. 2015. Improved white spruce (Picea glauca) genome assemblies and annotation of large gene families of conifer terpenoid and phenolic defense metabolism. The Plant Journal. 83:189–212. doi: 10.1111/tpj.12886.

Weber F, Dietrich A, Weil J-H, Maréchal-Drouard L. 1990. A potato mitochondrial isoleucine tRNA is coded for by a mitochondrial gene possessing a methionine anticodon. Nucleic Acids Research. 18:5027–5030. doi: 10.1093/nar/18.17.5027.

Wick RR, Judd LM, Gorrie CL, Holt KE. 2017a. Completing bacterial genome assemblies with multiplex MinION sequencing. Microbial Genomics. 3. doi: 10.1099/mgen.0.000132.

Wick RR, Judd LM, Gorrie CL, Holt KE. 2017b. Unicycler: Resolving bacterial genome assemblies from short and long sequencing reads Phillippy, AM, editor. PLOS Computational Biology. 13:e1005595. doi: 10.1371/journal.pcbi.1005595.

Wick RR, Schultz MB, Zobel J, Holt KE. 2015. Bandage: interactive visualization ofde novogenome assemblies: Fig. 1. Bioinformatics. 31:3350–3352. doi: 10.1093/bioinformatics/btv383.

Won H, Renner SS. 2003. Horizontal gene transfer from flowering plants to Gnetum. Proceedings of the National Academy of Sciences. 100:10824–10829. doi: 10.1073/pnas.1833775100.

